# *De Novo* Design, Directed Evolution and Computational Study of Heme-Binding Helical Bundle Protein Catalysts for Biocatalytic Enantioselective Ge–H Insertion

**DOI:** 10.1101/2025.09.29.679279

**Authors:** Wei Huang, Gessica M. Adornato, Maggie Horst, Turki M. Alturaifi, Kaipeng Hou, Peng Liu, William F. DeGrado, Yang Yang

**Affiliations:** Department of Chemistry and Biochemistry, University of California Santa Barbara, Santa Barbara, California 93106, USA; Department of Chemistry, University of Pittsburgh, Pittsburgh, Pennsylvania 15260, USA; Department of Pharmaceutical Chemistry & Cardiovascular Research Institute, University of California, San Francisco, California 94158, USA; Biomolecular Science and Engineering (BMSE) Program, University of California Santa Barbara, Santa Barbara, California 93106, USA

## Abstract

*De novo* designed proteins offer a malleable platform for the development of stereoselective transformations not found in biochemistry. Here, we report the *de novo* design and directed evolution of helical bundle protein catalysts for enantioselective germylation through Ge–H insertion, a transformation not previously achieved by enzymatic catalysis. Comparative computational analysis revealed that, relative to Si–H insertion, the Ge–H insertion reaction proceeds through an earlier and more flexible transition state, introducing distinct challenges for stereocontrol. Using a fully *de novo* designed truncated four-helix bundle scaffold as the starting point, directed evolution afforded a quadruple mutant that catalyzes Ge–H insertion with high efficiency, enantioselectivity, and broad substrate scope. Molecular dynamics simulations indicated that beneficial mutations introduced from directed evolution enhanced active-site preorganization and modulated local backbone flexibility, contributing to improved transition-state complementarity with fine-tuned binding pocket size and more stable cofactor positioning regulated by hydrogen bonding interactions. These findings showcase the excellent potential for *de novo* proteins to achieve stereoselective transformations previously unknown to biocatalysts and underscore the importance of active-site remodeling of *de novo* protein scaffolds via directed evolution in achieving selective catalysis involving flexible transition states.

**Table of Contents artwork:** 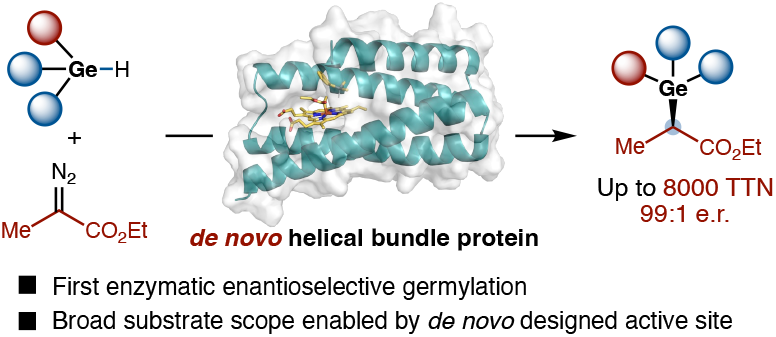

## Introduction

Due to their many attractive attributes, *de novo* designed protein catalysts hold the potential to significantly advance biological catalysis.^1^ Over the past decade, through the engineering of naturally occurring proteins, the biocatalysis community has developed a range of stereoselective catalytic reactions not previously found in nature.^2,3, 4^ Compared to natural proteins, *de novo* proteins can be designed to feature small molecular weight (<15 kDa) and excellent thermostability, making them excellent candidates for efficient and robust catalysts.^5^ Designed helical bundle proteins can also tolerate a variety of organic solvents,^6^ thus allowing hydrophobic substrates to be solubilized and transformed at higher concentrations (10^2^ mM) of relevance to larger-scale applications.^7^ Furthermore, through transition state-based protein design, *de novo* proteins can be rationally designed to furnish an excellent starting point for the development of stereoselective biocatalysts, allowing previously unknown non-native transformations to be realized in a rational manner within the cellular milieu.^8,9,10^

Recently, we designed metalloporphyrin binding four-helix bundle proteins as efficient biocatalysts for stereoselective cyclopropanation and silylation reactions which were not available in native enzymology (Figure 1(A)).^7^ As a key design consideration, we truncated the C-terminal helix of the four-helix bundle to craft an active site environment to selectively stabilize the preferred stereoisomeric transition states, thus allowing these carbene transfer reactions to proceed with excellent enantioselectivity. To design *de novo* heme-binding protein catalysts, we employed our previously developed bioinformatic tool, Convergent Motifs for Binding Sites (COMBS)^11^, to enhance cofactor-protein interactions, enabling the design of an enzyme with high enantioselectivity and total turnover number. Together, these results established truncated four-helix bundle *de novo* proteins as an excellent platform for the discovery and optimization of stereoselective new-to-nature reactions.

**Figure 1.**
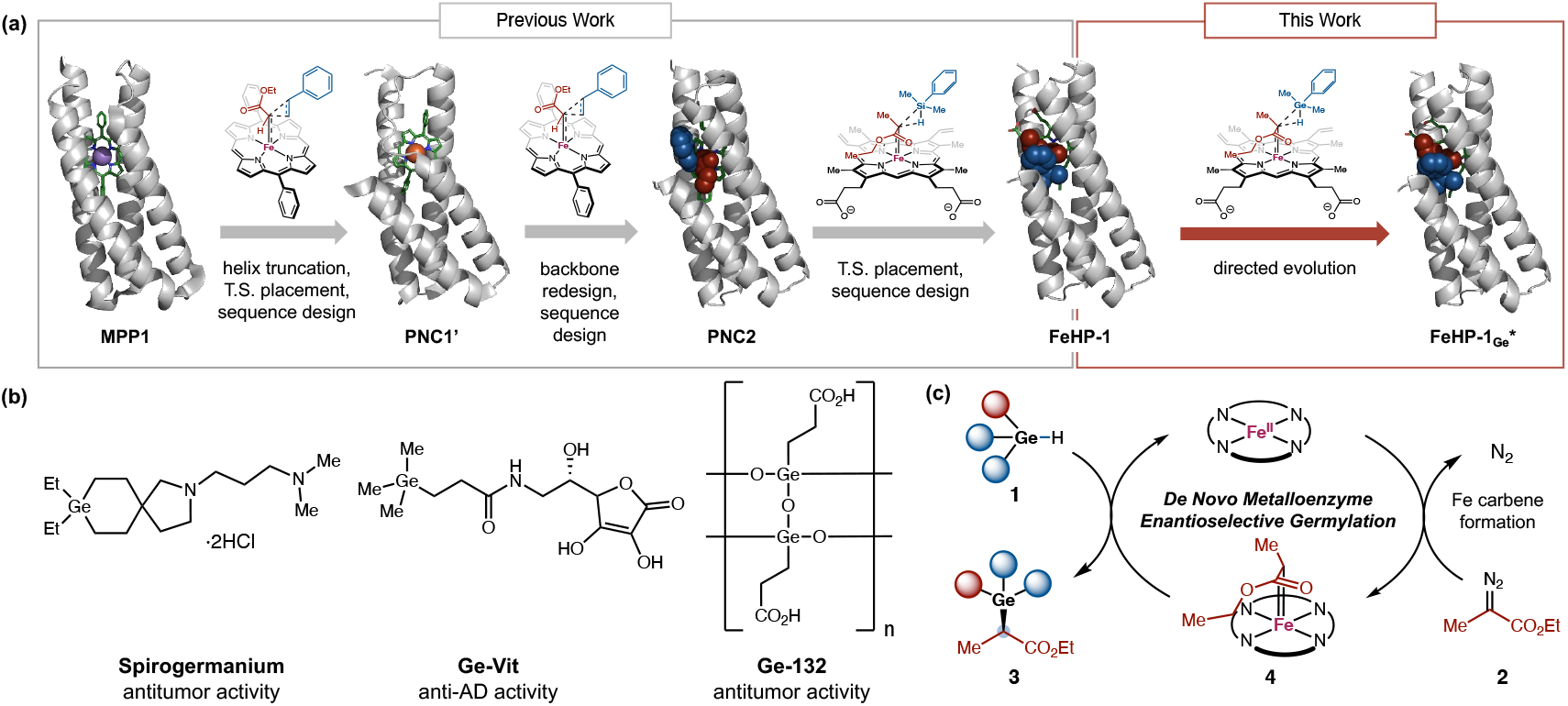
*De novo* Design and Directed Evolution of Heme-binding Four-Helix Bundle Protein Catalyst for Enantioselective Germylation. (a) Overall design and engineering strategy. Starting from MPP1 (PDB: 7JRQ), the C-terminal helix was truncated to accommodate the transition state (T.S.), and sequence design yielded PNC1’ (PDB: 9CTE). A loop deviation observed in the PNC1’ crystal structure was addressed by backbone redesign using MASTER^19^, followed by sequence design to generate PNC2, an efficient cyclopropanation catalyst. Placement of a Si–H insertion T.S. into PNC2 and subsequent sequence redesign led to the FeHP-1 scaffold. Directed evolution from FeHP-1 to FeHP-1_Ge_^*^ enabled enantioselective Ge–H insertion. (b) Representative bioactive germanium-containing molecules. (c) Proposed catalytic cycle for Fe-porphyrin-catalyzed enantioselective germylation via carbene insertion. Species 4 denotes the Fe-porphyrin–carbene intermediate. T.S. = transition state.

Based on these results, we asked whether we could further leverage this *de novo* protein platform to develop synthetically useful new-to-nature reactions not previously achieved with natural proteins. Previously, pioneering research by Arnold and Fasan demonstrated the use of engineered natural hemoproteins to catalyze Si–H and B–H insertion reactions.^12^ We turned our attention to biocatalytic asymmetric organogermane synthesis via Ge–H insertion reactions, a process not previously known with natural proteins. Due to the broad application of organogermanium compounds in materials science,^13^ synthetic chemistry,^14^ and medicine (Figure 1(B)), the stereoselective synthesis of organogermane is highly appealing.^15^ To date, relatively few methods have been established to allow for the catalytic asymmetric synthesis of organogermanium compounds.^16^ Among these, elegant examples of previously developed enantioselective Ge–H insertion reactions using small-molecule catalysts remain limited to the use of *α*-aryl substituted diazo compounds,^16a,16b,16d^ due to the preference of these transition-metal catalysts to confer enhanced enantioselectivities with donor-acceptor carbenes.^17^ In contrast, highly enantioselective Ge–H insertion reactions involving *α*-alkyl diazo substrates proved elusive.

We imagined this study as an opportunity to test the capacity of directed evolution^18^ to repurpose an enzyme designed *de novo* for one purpose, Si–H insertion reactions, to a new purpose, Ge–H insertion. This transformation forges new Ge–C and C–H bonds through two steps. In the first, highly exergonic step, iron catalyzes extrusion of molecular nitrogen from a diazo precursor to create a carbenoid species in which the methylene carbon interacts directly with Fe(II). In the second step, this carbenoid species inserts into a hydrogen-germanium bond, forming two new single bonds: one to H and one to Ge. The chirality at this carbon center is set by the geometry of the interaction between the metal carbene and the hydrogermane substrate. We expected that the energetic barrier for cleaving the Ge–H bond would be lower than that for cleaving the Si–H bond and the transition state would be more flexible, potentially leading to stereochemical promiscuity. Thus, achieving high enantioselectivity of the transformation remained a central challenge. If *de novo* heme protein catalysts could be designed and engineered to allow for enantioselective Ge–H insertion, it would constitute the first examples of biocatalytic C–Ge bond formation to address the unmet synthetic challenges with chiral small-molecule catalysts. Furthermore, the development of highly enantioselective germylation biocatalysts would provide an opportunity for the biosynthesis of organogermanium compounds, thereby bringing germanium, an uncharted inorganic element, to the periodic table of life.

### Results and discussion

To design and evolve helical bundle proteins for asymmetric Ge– H insertion, we selected dimethylphenylgermane (**1a**) and ethyl 2-diazopropionate (**2**, EDP) as model substrates (Figure 1(C)). At the outset of this study, we performed density functional theory (DFT) calculations to investigate the mechanism and reaction coordinate of the heme cofactor-catalyzed Ge–H insertion reaction. The Fepromoted N_2_ dissociation from EDP (**2**) forms a heme–carbene species **4**, which then undergoes carbene insertion into the Ge–H bond in **1a** to form the organogermane product (**3a**).^12d^ We computed the transition state (TS) for Ge–H insertion using a model heme system and compared this TS with that of previously studied Si–H insertion reactions (Figure 2).^7,12c,12d,20^ Similar to the Si–H insertion, the Ge– H insertion proceeds via a closed-shell singlet transition state (**TS1**), which is 6.2 kcal/mol more favorable than the triplet TS (Figure S1). The DFT-computed reaction energy profiles (Figure 2(A)) indicate that dimethylphenylgermane (**1a**) is significantly more reactive than dimethylphenylsilane (**1a′**) with a 2.0 kcal/mol lower kinetic barrier. Although both Ge–H and Si–H insertion transition states are concerted, leading to the corresponding organogermane and silane products directly without an intermediate, the Ge–H/Si–H bond cleavage is much more advanced than the Ge–C/Si–C bond formation in both transition states (Figure 2(B)). These structurally asynchronous transition state structures suggest that the reactivity of Ge–H/Si–H insertion is largely affected by the strength of the breaking Ge–H/Si–H bond.

**Figure 2.**
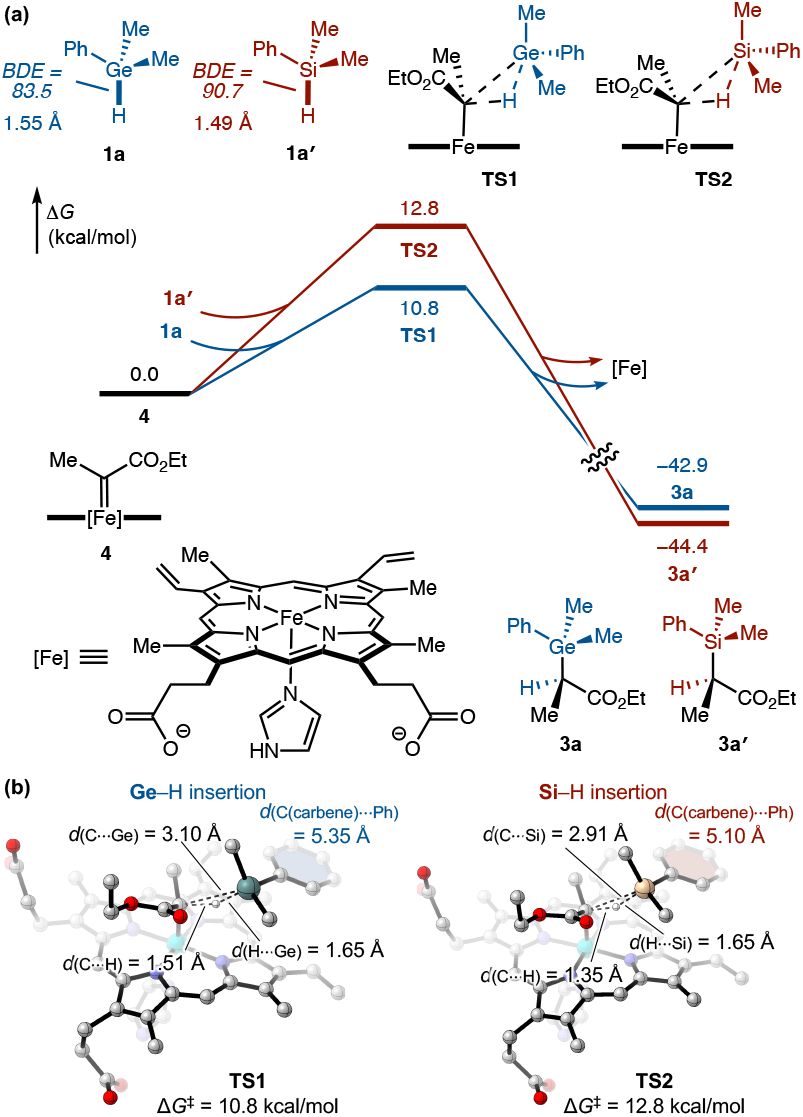
DFT calculations of the Ge–H and Si–H insertion reactions. (a) DFT-computed reaction energy profiles of Ge–H and Si–H insertion reactions. (b) Transition states of Ge–H and Si–H insertion.

The higher reactivity of **1a** can therefore be attributed to the Ge– H bond being weaker (BDE = 83.5 kcal/mol) than the Si–H bond in **1a′** (BDE = 90.7 kcal/mol). Though the van der Waals radii of Si and Ge are nearly the same, the bond length of Ge–H is longer than that of Si–H (Figure 2(A)). Considering that the Ge–H elongation from **1a** to **TS1** (0.10 Å) is smaller than the Si–H elongation from **1a′** to **TS2** (0.16 Å), the Ge–H insertion involves an earlier TS than Si–H insertion. This is also evidenced by the longer forming C– H/C–Ge bonds in **TS1** compared with those in the Si–H insertion **TS2**. Ge–H insertion’s earlier TS is consistent with the Hammond postulate.^21^ As a result of the earlier transition state and the slightly longer bond lengths to Ge than Si, the overall size of the Ge–H insertion TS is larger than the Si–H insertion TS by approximately 0.25 Å based on the distance between the carbenoid carbon and the benzene ring centroids (*d*_(C(carbene) ⋯Ph)_) in **TS1/TS2**. In addition, the Ge–H insertion TS is conformationally more flexible. The DFT-computed conformer ensembles, obtained from a conformational search workflow consisting of semiempirical tight-binding calculations followed by DFT refinement of the most stable conformers (see SI for details), revealed a greater number of low-energy TS conformers (Figure S2) and a reduced conformational preference of the germane substrate in the Ge–H insertion TS compared to the Si–H insertion TS.

Given the mechanistic similarity between Ge–H and Si–H insertion reactions, we postulated that the same set of designed *de novo* proteins could enable both enantioselective germylation and silylation reactions. Nonetheless, the differences in the size and flexibility of the DFT-computed transition states suggest that further enzyme engineering might be needed to account for the larger and more flexible Ge–H insertion TS. In addition, the higher reactivity of the hydrogermane substrate suggests it could be more difficult to achieve enzyme-controlled enantioselectivity compared to the corresponding Si–H insertion. Consequently, we thought the Ge–H insertion would be an ideal catalytic challenge through which to explore the capacity for computationally-guided directed evolution to refine enzymes composed by *de novo* design.

To assess whether our previously developed scaffolds could accommodate this more demanding reaction, we evaluated a set of ten *de novo* four-helix bundle proteins that bind heme. These scaffolds were derived from a parametrically generated D2-symmetric antiparallel bundle, SCRPZ-1, which was successively redesigned to improve folding stability (SCRPZ-2),^22^ to accommodate a hyper-stable folded core (PS1),^10d^ and to bind a manganese diphenyl porphyrin with axial histidine and water ligands (MPP1).^10e^ The helical bundle scaffold was then truncated and new loops were introduced between the helices to give the enzyme Porphyrin NovoChrome1 (PNC1), which showed high turnover frequency, turnover number, and enantioselectivity.^7^ To facilitate whole-cell screening, PNC1 was reengineered to bind heme by selecting favorable interactions for heme carboxylates using COMBS,^11a^ and the protein sequences were redesigned using LigandMPNN^23^ around the (*R*)-transition state of the Si–H insertion reaction. The top ten candidate scaffolds for Si–H insertion were selected based on evaluation of the degree to which the predicted structures accommodated the transition state structure, RaptorX^24^ pLDDT scores, and RMSD agreement between structures predicted by RaptorX and ESMfold^25^ (Figure 1(A)).

Initial screening revealed that all the *de novo* four-helix bundle proteins exhibited measurable catalytic activity, though most gave only modest yields or selectivities (Table S1). Among them, FeHP-1 exhibited the highest enantioselectivity, achieving an enantiomeric ratio (e.r.) of 87:13. Subsequently, we validated the performance of FeHP-1 in the form of whole-cell biocatalysts. It was found that for the current Ge–H insertion reaction, the use of whole cells or cell-free lysates of *de novo* proteins afforded similar results. At a biocatalyst loading of ca. 0.032 mol% (OD_600_ = 30), FeHP-1 catalyzed the Ge–H insertion of **1a** and **2** to afford product **3a** in 72% yield, 1880 ± 20 TTN, and 87:13 e.r., establishing a functional baseline for directed evolution (Table 1, entry 1). With FeHP-1 as the template, we next carried out directed evolution using site-saturation mutagenesis (SSM) and screening. In each SSM library generated by using the 22c-trick method,^26^ 88 clones were selected and screened using intact bacterial cells in a 96-well plate format. Guided by an AlphaFold3^27^ model of FeHP-1, we first targeted residue 70 proximal to the heme Fe center. In our previous study of *de novo* enzymes for Si–H insertion, we found that introduction of A70P substitution led to a slight kinking of the geometry of helix three that increased e.r. from 94:6 to 97:3 and increased TTN.^7^ In the current study, SSM of residue 70 and screening led to the discovery of FeHP-1 A70P as a superior germylation biocatalyst, improving the enantioselectivity of **3a** from 87:13 to 92:8 e.r. (entry 2). The introduction of an *α*-helix disrupting proline residue into H3 of the four-helix bundle represented a notable beneficial mutation that would be challenging to predict by the state-of-the-art protein design methods. We next targeted residue 106, which forms van der Waals contacts with the substrate fragment in the transition state. For the Si–H insertion reaction, we had previously found that the mutation F106V improved TTN and enantioselectivity. Given the role of this residue in shaping the hydrophobic interface between *α*-helices 3 and 4 and its proximity to the aryl group of the substrate, we hypothesized that substitutions at this site could impact the Ge–H insertion reaction. Indeed, SSM and screening at residue 106 led to FeHP-1A70P F106R, providing **3a** in 75% yield with 2360 ± 70 TTN and 97:3 e.r. (entry 3).

**Table 1.** Directed Evolution of FeHP-1 for Enantioselective Germanium-Hydrogen Insertion.

| entry | FeHP-1 variant | yield of **3a** | TTN of **3a** | e.r. of **3a** |
| --- | --- | --- | --- | --- |
| 1<sup>a</sup> | FeHP-1 | 60 ± 1 | 1,880 ± 20 | 87:13 |
| 2<sup>a</sup> | FeHP-1 A70P | 76 ± 1 | 2,390 ± 40 | 92:8 |
| 3<sup>a</sup> | FeHP-1 A70P F106R | 75 ± 2 | 2,360 ± 70 | 97:3 |
| 4<sup>a</sup> | FeHP-1 A70P F106R W63I S64E (FeHP-1<sub>Ge*</sub>) | 80 ± 3 | 2,500 ± 80 | 99:1 |
| 5<sup>b</sup> | FeHP-1<sub>Ge*</sub> | 94 ± 1 | 7,970 ± 80 | 99:1 |

<sup>a</sup>Reaction conditions: **1a** (5.0 mM, 1 equiv), **2** (7.5 mM, 1.5 equiv), 7.5% (v/v) MeCN in M9-N buffer. Reactions were carried out in triplicates using whole *E. coli* cells ( $OD_{600}$ = ca. 30) overexpressing FeHP-1 variant (ca. 0.032 mol%). Heme loading was determined by the hemochrome assay as detailed above. Averaged yields and TTNs are shown. <sup>b</sup>Reaction conditions: **1a** (5.0 mM, 1 equiv), **2** (7.5 mM, 1.5 equiv), 20% (v/v) EtOH in M9-N buffer. Reactions were carried out in triplicates using whole *E. coli* cells ( $OD_{600}$ = 15) overexpressing FeHP-1 A70P F106R W63I S64E (ca. 0.0118 mol%).

To further enhance stereocontrol, we turned to the H2–H3 loop, a flexible segment near the active site. The H2–H3 loop was initially introduced via relooping using a previously developed loop search program, MASTER^19^, to replace a distorted portion of helix 3 that clashed with the cofactor as revealed by previous X-ray crystallography.^7^ Given its proximity to the substrate binding pocket, we hypothesized that refining its sequence could alter loop rigidity and influence the overall protein dynamics, thus affecting the catalytic activity and enantioselectivity of the *de novo* protein. Single-site saturation mutagenesis and screening targeting residues 62–64 identified beneficial substitutions at positions 63 and 64 (Table S3). To explore the potential epistatic effect, we created a double site saturation library by simultaneously randomizing residues 63 and 64. Screening of a total of 400 clones revealed that a double mutant W63I S64E improved the enantioselectivity from 97:3 1 e.r. to 99:1 e.r. and the yield from 75% to 80% (entry 4). Further site-saturation mutagenesis at C-terminal residues 112 and 106 did not result in improved protein variants. Thus, the final evolved variant, FeHP-1 A70P F106R W63I S64E (FeHP-1_Ge_^*^), combined high catalytic activity and enantioselectivity, affording the Ge–H insertion product **3a** in 80% yield and 99:1 e.r. (entry 4).

Overall, these results reveal that a single *de novo* designed protein can be successfully adapted to catalyze two different new-to-nature reactions in high enantioselectivity through directed evolution. The evolutionary landscape traversed by the two design campaigns is shown in Figure 3. The reactivity for the Ge–H insertion reaction is consistently higher, as expected based on its increased reactivity. However, the key problem of enantioselectivity was more challenging for this reaction than for Si–H insertion. By making multiple mutations in the loop region, this directed evolution strategy yielded an enzyme with enantioselectivity for Ge–H insertion that surpassed its performance for Si–H insertion and nonetheless delivered excellent yield.

**Figure 3.**
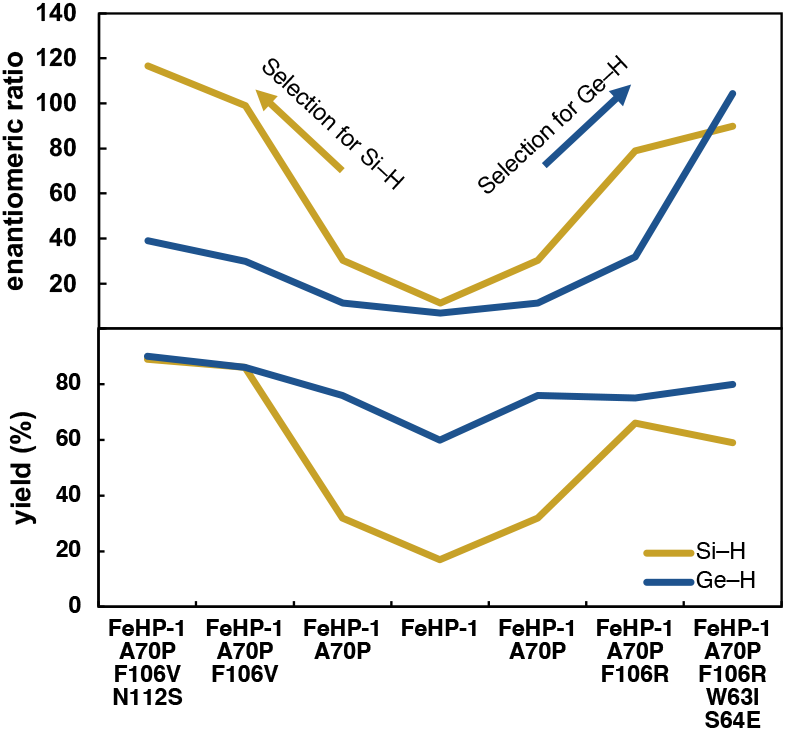
Enhancement of enzyme properties during the two directed evolution campaigns. Enantiomeric ratio (e.r.) is plotted as (*R*)/(*S*). Reactions were carried out in whole cells for 2 h (OD_600_ = ca. 10 for Si– H and ca. 30 for Ge–H).

Further reaction condition optimizations revealed ethanol (EtOH) as the optimal co-solvent for whole-cell biotransformations (Table S5). The evolved protein catalyst FeHP-1_Ge_^*^retained excellent activity, and enantioselectivity was found to tolerate up to 20% (v/v) EtOH in whole-cell. Optimal results were obtained with 20% (v/v) EtOH using intact *E. coli* cells (OD_*600*_ = 15). Using these conditions, FeHP-1_Ge_^*^afforded product **3a** in 94% yield, 7970 ± 80 TTN, and 99:1 e.r. (Table 1, entry 5). Furthermore, FeHP-1_Ge_^*^retained high stability to high concentrations of EtOH, a desirable green organic solvent that effectively solubilizes common organic substrates (Figure 4(A) and Table S6). With FeHP-1_Ge_^*^, the yield and enantioselectivity of the Ge–H insertion product **3a** remained unchanged with up to 40% (v/v) EtOH in water, demonstrating the exceptional stability of *de novo* designed protein catalysts in the presence of organic solvents. The excellent organic solvent compatibility of our *de novo* proteins in turn allowed the biocatalytic reactions to occur with substantially increased substrate concentrations (Figure 4(B)). Using 30% EtOH as the solvent, the yield and e.r. of product **3a** remained almost unaffected with **1a** concentration of up to 100 mM, about 10-fold higher than related new-to-nature carbene transfer reactions catalyzed by engineered natural heme proteins^12a,12e^ (see Table S9 for further details).

**Figure 4.**
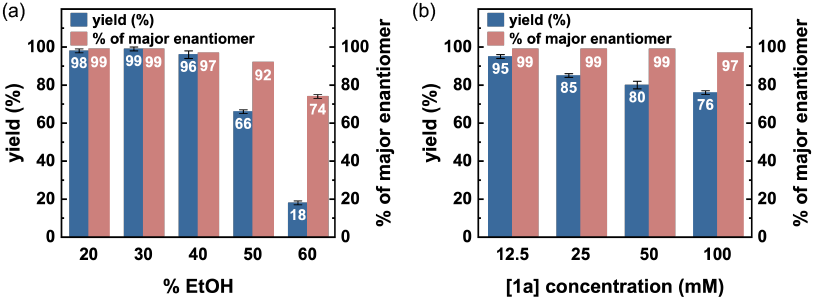
Reaction Results of High Percentage of EtOH and High Substrate Concentration. (a) Biocatalytic Ge–H insertion with different amounts of EtOH. Reaction conditions: **1a** (5.0 mM, 1 equiv), **2** (7.5 mM, 1.5 equiv), EtOH in M9-N buffer. Reactions were carried out in triplicates using cell-free lysate (OD_600_ = ca. 15) overexpressing FeHP-1_Ge_^*^ (ca. 0.026 mol%). (b) Biocatalytic Ge-H insertion with different substate concentration of hydrogermane substrate. Reaction conditions: **1a** (1 equiv), **2** (1.5 equiv), 30% (v/v) EtOH in M9-N buffer. Reactions were carried out in triplicates using purified FeHP-1_Ge_^*^ (0.01 mol% loading).

With the newly evolved Ge–H insertion biocatalyst FeHP-1_Ge_^*^, we examined its substrate scope under the optimized reaction conditions (Table 2). Aryl-substituted germane derivatives bearing a substituent at the *ortho*-(**3b**), *para*-(**3d**), and *meta*-positions (**3c**) were well tolerated, affording the corresponding Ge–H insertion products in excellent yields and enantioselectivities. A variety of electron-donating groups at the *para*-position, including a methoxy (**3e**), a dimethylamino (**3f**), and a hydroxy (**3g**) group, were compatible, allowing a range of enantioenriched organogermanium products to be prepared with excellent yields. The compatibility of sensitive functional groups such as phenol underscored the evolved catalyst’s functional group tolerance and chemoselectivity, favoring Ge–H insertion over O–H insertion. In addition, an electronically neutral methylthio group (**3h**) and halogen substituents including a fluoro (**3i**) and a chloro (**3j**) group were readily accommodated. Furthermore, in contrast to the previously studied Si–H insertion^12a^, an electron-deficient aryl group possessing a 4-trifluoromethyl (**3k**) was also compatible with this Ge–H insertion reaction, further showcasing the excellent substrate tolerance. Furthermore, substrates with a bulky substituent such as a biphenyl (**3l**), a 2-naphthyl (**3m**) and a benzofuran (**3n**) could be effectively transformed, suggesting that the truncated four-helix bundle protein is insensitive to the steric properties of the substrate, a feature that is uncommon with natural proteins with a confined active site. Finally, phenyldiethylgermane could also be applied to afford the desired chiral organogermanium product (**3o**) with excellent yield and enantioselectivity, further underscoring the broad substrate tolerance of FeHP-1_Ge_^*^.

**Table 2.**
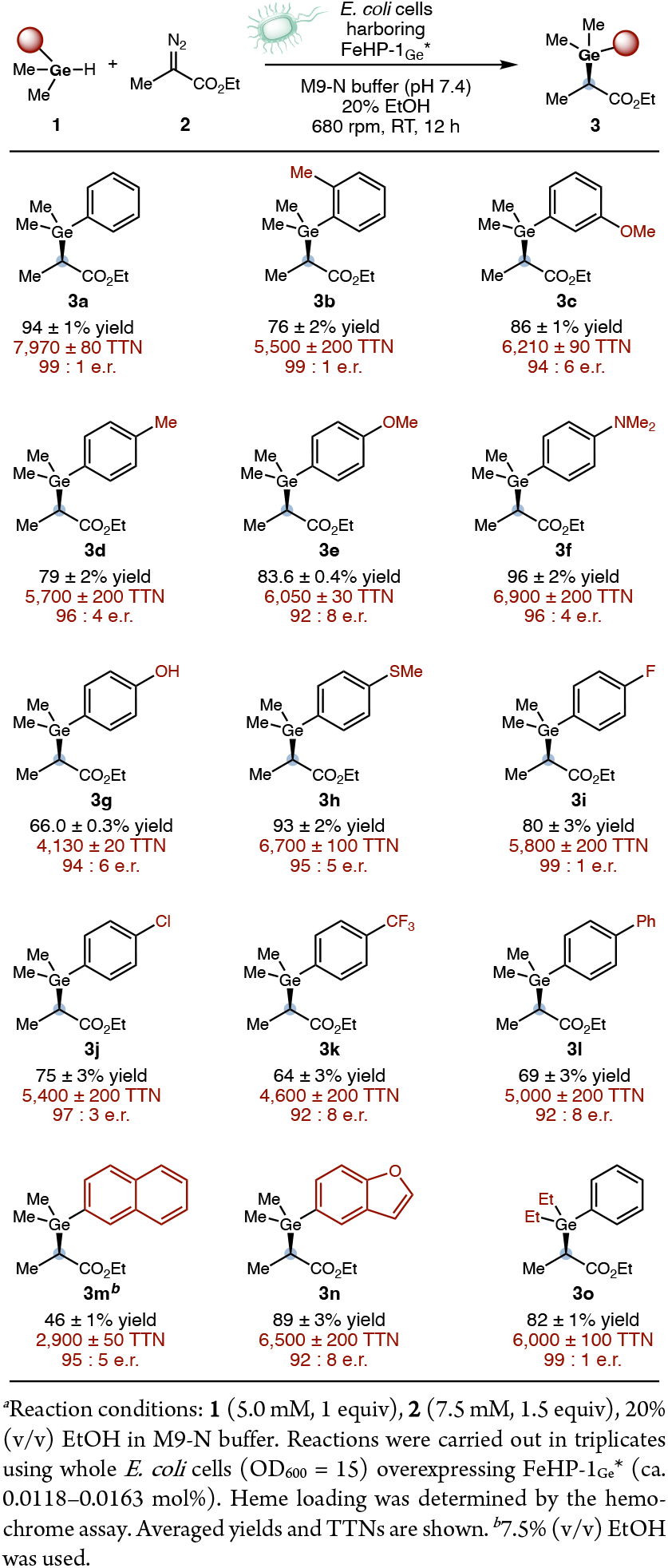
Substrate Scope of Biocatalytic Enantioselective Ge-H Insertion Using Evolved *De novo* Protein FeHP-1_Ge_^\**a*^.

To unambiguously establish the absolute configuration of the Ge–H insertion products, we converted ester **3a** (99:1 e.r.) to the corresponding amide **6** via amidation with 4-bromoaniline (**5**), following a reported procedure.^28^ The resulting amide **6** retained good levels of enantiopurity (95:5 e.r.). X-ray single crystal diffraction analysis of **6** confirmed its absolute configuration to be (*R*)-, consistent with the stereochemical model of our transition state-based design (Figure 5).

**Figure 5.**
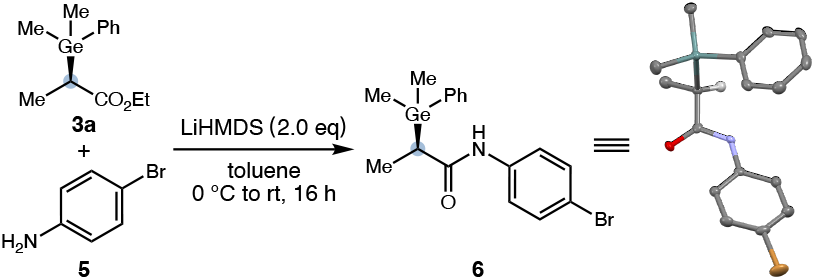
Derivatization and Determination of Absolute Stereochemistry of the Ge–H Insertion Product by X-Ray Diffraction Analysis. Thermal ellipsoids were set at 50% probability; hydrogen atoms except the one attached to the stereogenic carbon were omitted for clarity.

To better understand how beneficial mutations uncovered by directed evolution affect enzyme structure and catalytic efficiency, we performed classical molecular dynamics (MD) simulations of the best-performing variants from each round of directed evolution: FeHP-1, FeHP-1 A70P, FeHP-1 A70P F106R, and FeHP-1_Ge_^*^, as well as our previously reported best-performing variant for the Si–H insertion reaction, FeHP-1_Si_^*^ (FeHP-1 A70P F106V N112S). For each variant, two different types of simulations were performed, one involving the heme–carbene complex derived from EDP, and the other representing the near-attack conformation (NAC) of the heme–carbene with the hydrogermane substrate 1a. In the NAC simulations, the Ge atom of 1a was restrained at an average distance of 3.5 Å from the carbenoid carbon and placed at the (*Re*)-face of the carbene, leading to the major enantiomer of the product 3a (see SI for simulation setup details)

The heme–carbene simulations revealed that the active site preorganization improved following the introduction of beneficial mutations during the first two rounds of directed evolution (A70P and F106R). Figure 6(A) shows the distance between helices 3 and 4, the two helices that form the substrate binding pocket, for FeHP-1, FeHP-1_Si_^*^, and FeHP-1_Ge_^*^. In the parent protein (FeHP-1), two distinct distance populations are observed, centered around 9.5 and 11 Å, respectively, reflecting binding pocket heterogeneity due to variable helix–helix association. In FeHP-1_Si_^*^, the shorter H3–H4 distance population becomes the overwhelming majority, indicating a much tighter binding pocket with strong association between H3 and H4. FeHP-1_Ge_^*^, by contrast, continues to sample both conformations. This suggests that the mutations in FeHP-1_Ge_^*^ confer only a modest increase in active-site preorganization relative to FeHP-1, resulting in a more flexible and larger binding pocket compared to FeHP-1_Si_^*^. This difference likely reflects the need for FeHP-1_Ge_^*^ to accommodate and stabilize a larger and more flexible transition state associated with Ge–H insertion (*vide supra*). Consistent with this, NAC simulations show that the average distance between H3 and H4 in the NAC of FeHP-1_Ge_^*^ is 0.4 Å longer than that in FeHP-1_Si_^*^ (Figure 6(C)). The longer H3–H4 distance in FeHP-1_Ge_^*^ creates a larger cavity to accommodate the ca. 0.25 Å larger transition state for Ge–H insertion compared to Si–H insertion (Figure 2(B)).

**Figure 6.**
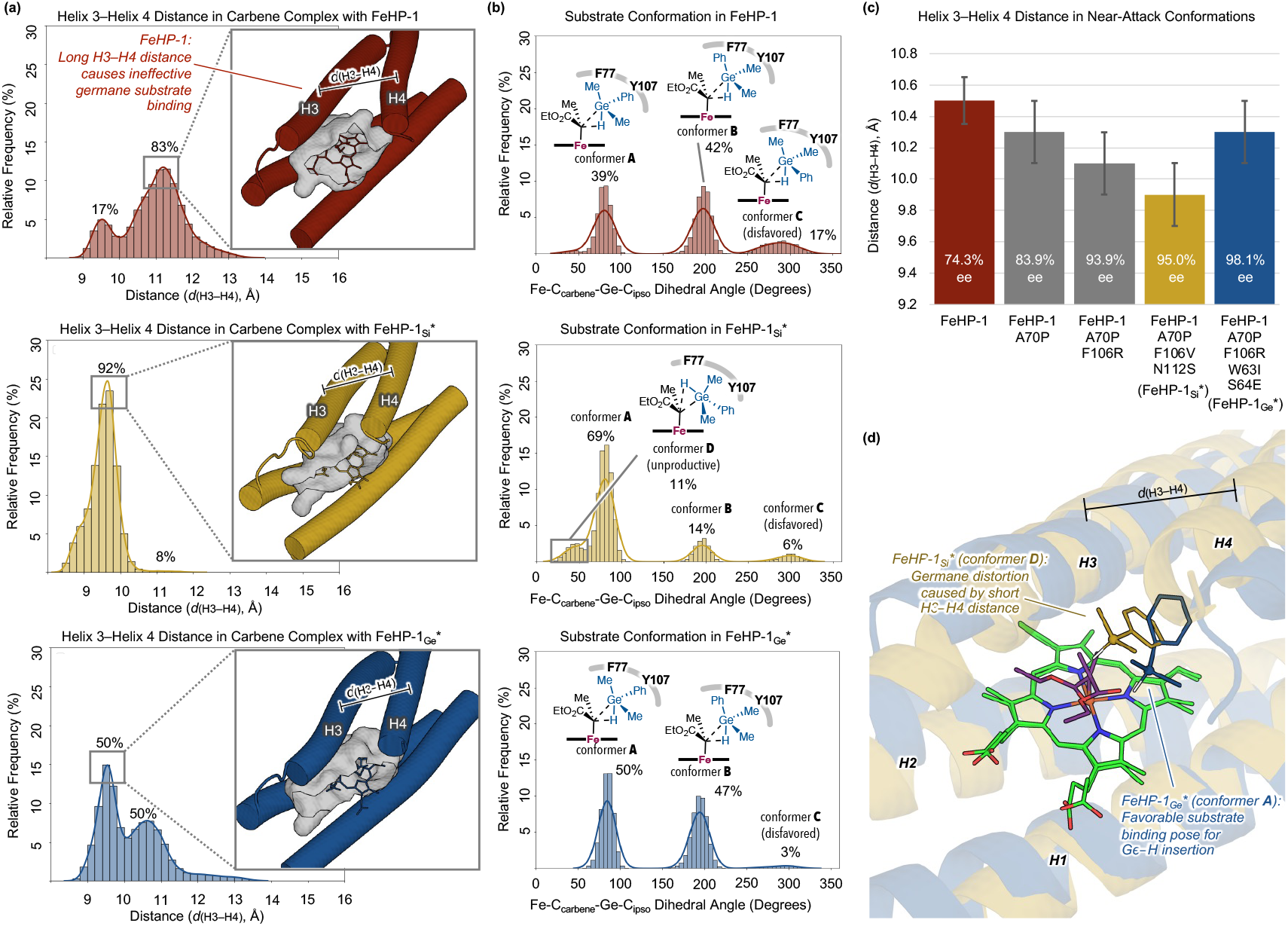
Effects of binding pocket size on substrate conformation and active-site organization. (a) Distance between helices 3 (H3) and 4 (H4) in the heme–carbene complexes of FeHP-1, FeHP-1_Si_^*^, and FeHP-1_Ge_^*^. Pocket cavity surfaces are shown for each variant. Distances were calculated as the separation between the centroids of the backbone heavy atoms in H3 (residues 67–89) and H4 (residues 94–112) at 2 fs intervals throughout the MD simulations. (b) Conformations of the hydrogermane substrate in NAC simulations. (c) Average H3–H4 distances in NAC simulations of each variant. Experimentally determined enantiomeric excess (*ee*) values for the Ge–H insertion reaction are displayed on each bar. (d) Representative structure of conformer **D** from the NAC simulation of FeHP-1_Si_^*^ (gold) showing distorted hydrogermane substrate geometry, overlaid with a representative conformer **A** structure from FeHP-1_Ge_^*^ (blue). H1 = helix 1, H2 = helix 2, H3 = helix 3, H4 = helix 4.

Next, we analyzed the conformation of the hydrogermane substrate in the NAC simulations to examine how the mutations influence the substrate binding pose and geometrical distortion required to reach the Ge–H insertion transition state (Figure 6(B)). In the NAC with the parent FeHP-1, the hydrogermane substrate could adopt three distinct conformations (conformers **A–C**). In conformers **A** and **B**, the phenyl group of the substrate is nestled within the hydrophobic pocket surrounded by F77 and Y107. By contrast, conformer **C** adopts a syn-periplanar orientation relative to the ester group of the carbene, positioning the phenyl group toward the solvent. Our DFT calculations indicate that conformers **A** and **B** correspond to the lowest-energy transition states for Ge–H insertion, with conformer **B** only 0.7 kcal/mol higher in energy than conformer **A** (Figure S3). Ge–H insertion via conformer **C**, however, is predicted by DFT to be significantly less favorable, with a barrier of 2.1 kcal/mol higher than that of conformer A. The relatively high population of the nonproductive conformer C observed in FeHP-1 NAC simulations suggests that the active site in the parent *de novo* protein does not consistently enforce a reactive substrate geometry toward Ge–H insertion.

By contrast, the evolved FeHP-1_Ge_^*^ variant exhibited a substantially reduced population of conformer C in the NAC simulations. This is likely due to improved preorganization of the active site pocket, which stabilizes conformers A and B through more defined hydrophobic interactions between the phenyl group and residues lining the pocket between helices 3 and 4. In NAC simulations of FeHP-1_Si_^*^, which possesses the most compact active site pocket, conformer A is the dominant binding mode, and conformer C is similarly suppressed. However, a new binding pose (conformer D) emerges in these simulations, characterized by the Ge–H bond pointing away from the heme Fe, rendering the conformation unproductive. This geometry was not observed in the DFT-optimized conformer ensemble of the Ge–H insertion TS. We attribute this distortion to the limited pocket volume in FeHP-1_Si_^*^: the short distance between H3 and H4 restricts accommodation of the phenyl ring of the substrate 1a, forcing it out of the hydrophobic pocket and thereby distorting the substrate conformation (Figure 6(D)).

Next, we investigated the roles of two beneficial mutations, W63I and S64E, located in the loop connecting helices 2 and 3. The root mean square fluctuation (RMSF) analysis revealed that this loop is more flexible in FeHP-1 A70P F106R than in FeHP-1_Ge_^*^, as indicated by the higher RMSF values in the former variant (Figure 7(A)). Consistent with this, clustering analysis (Figure 7(B)) shows that the loop adopts a more disordered ensemble of conformations in FeHP-1 A70P F106R compared to the more rigid and ordered loop observed in FeHP-1_Ge_^*^. This difference in loop dynamics significantly impacts the hydrogen bonding interactions that anchor the heme cofactor. In FeHP-1_Ge_^*^, the rigidified loop allows the heme-6-propionate to form persistent hydrogen bonds with R7 and S67 across the majority of the MD frames (Figure 7(C)). By contrast, the increased backbone fluctuation in FeHP-1 A70P F106R leads to more dynamic and less consistent hydrogen bonding at the heme-6-propionate site, with additional hydrogen bonding interactions observed with S57, S64, and W63. By mutating residues 63 and 64 to non-hydrogen-bond-donor residues (W63E and S64I, respectively), FeHP-1_Ge_^*^ achieves a more well-defined hydrogen bonding network that stabilizes the heme cofactor in a consistent position. This anchoring mode closely resembles that observed in previous MD simulations of the Si–H insertion protein catalyst.^7^

**Figure 7.**
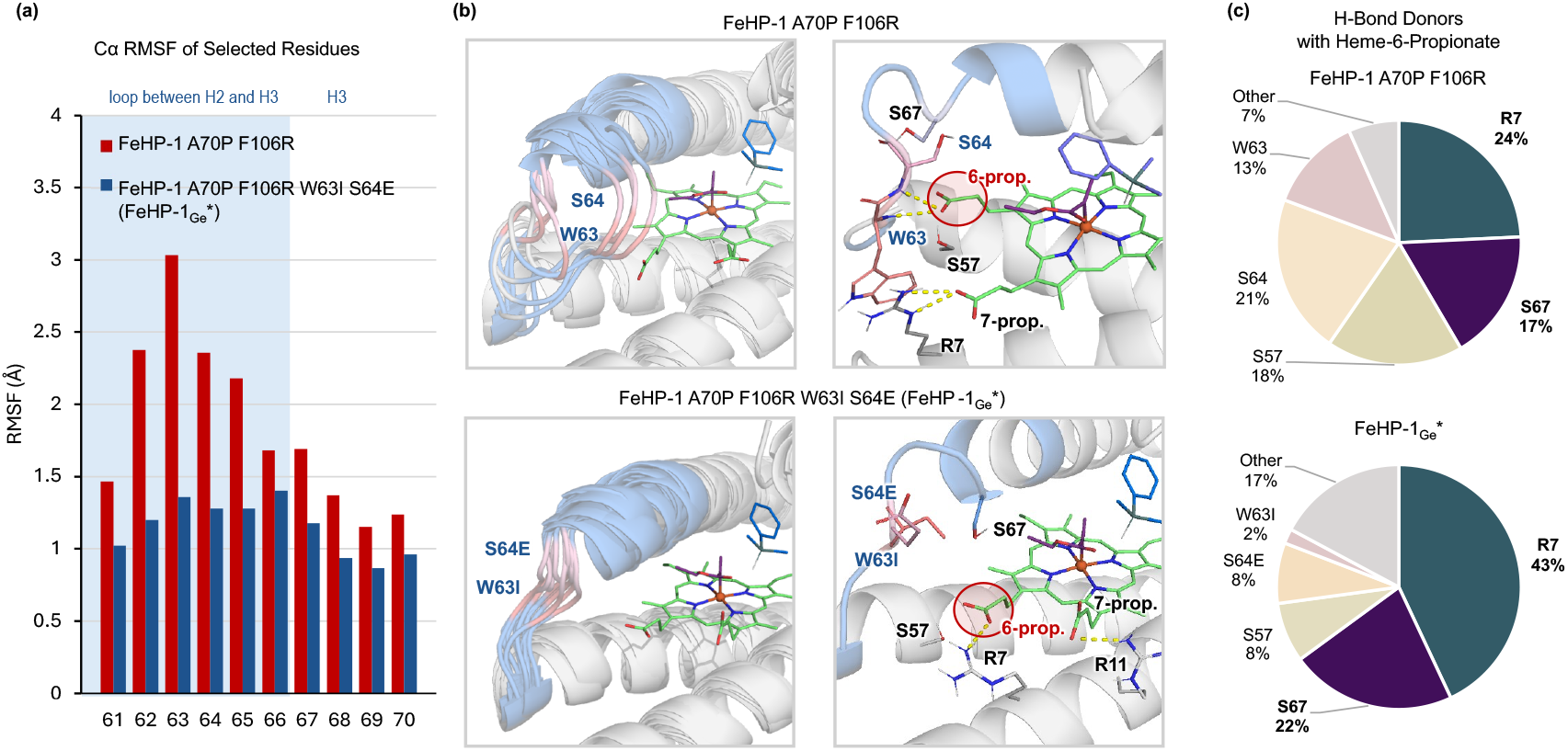
Effects of loop dynamics on heme cofactor binding. (a) RMSF values for the C*α* atoms of selected residues in the loop region and helix 3 in NAC simulations of FeHP-1 A70P F106R and FeHP-1_Ge_^*^. The loop region, corresponding to residues 61–66, is shaded in blue, while helix 3 is in light grey. (b) An overlay of representative structures based on clustering the backbone heavy atoms of residues 61–70 in FeHP-1 A70P F106R and FeHP-1_Ge_^*^. Representative structures demonstrating the hydrogen bonding with the heme propionates are shown. (c) The percentage of frames where hydrogen bonds are present between heme-6-propionate and residues near the active site. Water-bridged hydrogen bonds are included in each category (see Figure S16 for detailed analysis).

## Conclusion

In summary, we have demonstrated that *de novo* design and directed evolution can be effectively combined to generate heme-binding helical bundle proteins capable of catalyzing enantioselective Ge–H insertion, a new-to-nature transformation not previously achieved using biocatalysts. Starting from a four-helix bundle protein with a truncated C-terminus and a stabilized H2-H3 loop, transition-state-based design furnished 10 proteins for carbene insertion into Si–H bonds. Through three rounds of directed evolution applied to a *de novo* protein scaffold, we obtained a quadruple mutant, FeHP-1_Ge_^*^, with improved catalytic efficiency, enantioselectivity, and substrate compatibility. Computational analysis revealed that the beneficial mutations enhance active-site preorganization, modulate loop dynamics, and reestablish hydrogen-bonding interactions with the heme cofactor. These changes help accommodate the early and flexible transition state of the Ge–H insertion, thereby enabling effective stereocontrol. Taken together, these findings highlight the power of combining *de novo* protein design with laboratory directed evolution and computational analysis to expand the reach of protein catalysis to reactions not previously known in nature. More broadly, these studies offer a general strategy for enabling previously unavailable new-to-nature reactions in protein catalysis.

## ASSOCIATED CONTENT

### Supporting Information

The Supporting Information is available free of charge on the ACS Publications website.

Experimental procedures, DNA and protein sequences, characterization data, results of single-crystal X-ray analysis, computational details, HPLC traces, and NMR spectra (PDF)

Crystallographic data for compound (*R*)-6 (CIF)

Deposition number 2444539 (for (*R*)-6) contains the supplementary crystallographic data for this paper. These data are provided free of charge by the joint Cambridge Crystallographic Data Centre and Fachinformationszentrum Karlsruhe <u>Access Structures</u> service.

## ACKNOWLEDGMENT

*De novo* protein design and directed evolution studies are supported by the National Science Foundation (ITE-2448848 to Y.Y. and W.F.D.). Computational studies are supported by the National Science Foundation (CHE-2400087 to P.L. and CHE-2108660 to W.F.D.). Y.Y. is an Alfred P. Sloan Research Fellow (FG-2024-22244), a Camille Dreyfus Teacher-Scholar Awardee (TC-25-084) and a David & Lucile Packard Fellow (2023-76169). M.H. is supported by an Institutional Research Service Award in the Molecular Cellular Basis of Cardiovascular Diseases from the National Institutes of Health (T32HL007731-31A1). is supported by the National Institutes of Health (R35GM122603). The content is solely the responsibility of the authors and does not necessarily represent the official views of the National Institutes of Health. Computational studies were carried out at the University of Pittsburgh Center for Research Computing and Data and the Advanced Cyberinfrastructure Coordination Ecosystem: Services & Support (ACCESS) program, supported by NSF award numbers OAC-2117681, OAC-1928147, and OAC-1928224.

